# Unsupervised Identification of Cancer Attractor States through the Lens of Embryonic Origin and Cancer Hallmarks

**DOI:** 10.64898/2026.01.13.699215

**Authors:** Prashant Deshpande, Prabal Deb, Ibrahim Al Haddabi, Boris Itkin

**Affiliations:** Department of Clinical Laboratories, Sultan Qaboos Comprehensive Cancer Care and Research Center, University Medical City, Muscat, Oman

## Abstract

**Background:** Oncogenesis is a highly intricate process characterized by the transition of normal cells into aberrant biological attractor states. These states emerge from the non-random combinatorial interactions between inherent lineage-specific programs, governed by Gene Regulatory Networks (GRNs), and acquired somatic mutations. While the hallmarks of cancer provide a functional framework for transformation, current genomic models often fail to account for the topographical constraints of the cell of origin. We hypothesized that integrating embryological origin (EO) with hallmark-related mutations (HRM) enables the unsupervised identification of stable, pan-cancer attractor states.

**Methods:** Utilizing the MSK-MET database (n=25,775), we annotated somatic variants across 15 hallmarks. We developed and compared two unsupervised clustering models using Jaccard-distanced t-SNE and K-means algorithms: one based solely on HRM profiles and another integrating EO with HRM (EO/HRM). Model robustness and attractor stability were evaluated via bootstrap resampling and five-fold cross-validation.

**Results:** Hallmark-related mutations were identified in 95.5% of tumors. The EO/HRM model identified 11 distinct clusters and demonstrated superior stability (ARI: 0.74 vs. 0.70) and prognostic performance (C-index: 0.59 vs. 0.56) compared to the HRM-only model. These clusters represent stable functional basins, such as the TP53-driven minimalist attractor in Cluster C6 and the hyper-unstable attractor in Cluster C10. The clusters exhibited unique, organ-specific metastatic predilections independent of histological subtypes. Critically, we observed prognostic inversions, such as Cluster C0 representing a favorable attractor in lung adenocarcinoma but an aggressive state in prostate carcinoma, illustrating that attractor behavior is gated by the embryological landscape.

**Conclusion:** Integrating embryological origin as a proxy for inherent cellular programming significantly enhances the identification of stable, prognostically relevant biological attractor states. This framework provides a novel strategy for functional risk stratification and the development of context-aware precision therapies aimed at destabilizing malignant attractors.

## Introduction

Oncogenesis is an intricate, and highly heterogeneous process. Traditionally, this is conceptualized as a serial acquisition of somatic genomic alterations resulting in transformation of normal cells into neoplastic cells with abnormal phenotype and novel functional properties like autonomous proliferation, dedifferentiation, and tissue invasion(1). However, the observation that many disparate malignancies share common genomic variants yet exhibit significantly different clinical behaviors suggests that sequence data alone cannot fully explain the behavior of neoplastic cells.

The modern interpretation of Waddington’s epigenetic landscape and canalization frames the normal cell as a dynamic attractor state, where the natural history and behavior of cells is governed by an inherent program mediated through Gene Regulatory Networks (GRNs). Based on this understanding, cancer cells can be conceptualized as evolving aberrant attractor states emerging from non-random combinatorial interactions between altered internal program (GRN) and somatic sequence variants(2,3). However, testing this hypothesis in real life does pose significant challenges.

While the mechanisms of common individual gene variants are well characterized with respect to cellular pathways or mechanisms (e.g.,RAS/ RAF pathway or mismatch repair pathway), their ‘functional’ contribution to oncogenesis is less established(4,5). This is further complicated by pleiotropy; where a single mutation influences diverse cellular processes, and conversely, alterations in multiple genes may converge on shared cellular pathways. The “hallmarks of cancer” provides a functional framework for understanding the core biological processes enabling malignant transformation and progression. (6,7) By mapping genes to associated cancer hallmarks, a genetic variant can be interpreted in terms of functional impact on oncogenesis(8). Thus, somatic variants can be defined as drivers of oncogenesis via modulation of pre-existing programs and pathways, either enhancing or attenuating their intrinsic functions or ectopically activate otherwise dormant or lineage-inappropriate properties. These modulated or acquired properties are best captured by the framework of cancer hallmarks.

Furthermore, determining the inherent cellular program of organ-specific tumor-initiating cells remains elusive. Emerging evidence suggests that these cells resemble embryonic progenitors with respect to gene expression profiles and cellular properties (like proliferation, tissue invasion) with transcriptional program in carcinomas showing more resemblance to embryonic stem cells than adult stem cells(9). These similarities support the hypothesis that tumor-initiating cells retain or re-acquire functional programs reminiscent of the embryonic cells responsible for development of affected organs(9–12).

In line with this approach, we hypothesize that the embryonic origin (EO) for the involved organ can serve as a viable proxy for the inherent cellular programs (genetic regulatory network), while somatic genetic mutations—categorized based on their association with specific cancer hallmarks (hallmark related mutations or HRM)—can represent the modulated or acquired cellular properties. This hypothesis was tested using models based on HRM profile and embryonic cell of origin (EO/HRM) and HRM profile only.

## Methods

### MSK-MET Database and Variant Analysis

For study, we employed a large-scale publicly available MSK-MET database.(13) The relevant files were downloaded from cBioportal and a composite database was generated. All single nucleotide variants (SNVs), insertions/deletions (indels) and structural variants (translocations, deletion, inversion) resulting in a protein fusion were incorporated in a variant database. For translocation/fusion variants, both partner genes were incorporated into the analysis to capture the full spectrum of genomic alterations. Other structural variants and copy number variations (CNVs) were not included in this analysis due to lack of clarity regarding interpretation of variants and assessment of the impact of these variants.

### Definitions and Source Databases

The hallmarks refer to biological capabilities acquired by cells during the development of cancer and the following hallmarks of cancer were included in the analysis : invasion and metastasis (invasion/metastasis), evading growth suppressors (growth suppression), proliferative signaling (signaling), angiogenesis, differentiation and development (dysdifferentiation), global regulation of gene expression (genomic regulation), change of cellular energetics (energetics), genome instability and mutations (genomic instability), escaping programmed cell death (cell death), cell division control (cell division), tumour promoting inflammation (inflammation), cell replicative immortality (replicative immortality), senescence, escaping immune response to cancer (immune response), interaction with pathogen (pathogen response)(6).

Genes associated with individual hallmarks were extracted from the COSMIC (Catalogue of Somatic Mutations in Cancer) database (last accessed July 2025), and all variants were annotated individually. (14,15) A hallmark was considered involved if at least one variant was identified in any of the hallmark associated genes. Since a single gene can have multiple hallmark associations, a single variant in such gene would be annotated as involvement of more than one hallmark. In view of high representation of *TP53* gene across multiple hallmarks (13/15 hallmarks as described in COSMIC database), presence of TP53 variants was defined as a separate group. A final list of cancer hallmarks and associated genes is enlisted in supplementary table S1.

For the embryological origin of the involved organs, we identified the following subgroups: ectoderm, mesoderm, endoderm, neural crest, and yolk sac. The endoderm was further subdivided into foregut, midgut, hindgut, and urogenital sinus. A list of cancer types and corresponding embryological origins is described in the supplementary table S2.

### Model development

For analysis, hallmark involvement was coded as binary data (involved vs. not involved), based on the presence or absence of variant(s) in any one of the hallmark-associated genes. Embryological origins were represented as binary indicator variables via one-hot encoding, with each distinct origin assigned to an individual column. The MSK-MET database was divided into three subsets: training (n= 16496) validation (n = 4124) and test databases. The training and validation dataset (n=20620, 80%) were used for pattern identification, classifier development, while test dataset (n = 5,155, 20%) was used for confirmation of model performance and prognostic implications. To map the high-dimensional genomic-lineage landscape into a biologically interpretable space, we utilized t-Distributed Stochastic Neighbor Embedding (t-SNE). This nonlinear dimensionality reduction technique is particularly effective at preserving local structures in complex datasets. Given the binary nature of our hallmark-related mutation (HRM) and embryological origin (EO) data, the **Jaccard distance metric** was employed. Unlike Euclidean distance, which can be sensitive to “joint absences” in sparse binary vectors, the Jaccard metric focuses specifically on shared positive features, providing a more accurate representation of biological similarity between tumors(16). The two-dimensional coordinates generated by t-SNE served as the input for K-means clustering. This allowed for the unsupervised identification of discrete biological “attractor states” within the EO/HRM landscape. The optimal number of clusters was determined using the **elbow method** (Within-Cluster Sum of Squares) combined with a qualitative assessment of cluster separation. Because the high-dimensional complexity of the latent space made manual assignment of new cases unfeasible, we trained Random Forest classifiers on the training subset to automate cluster assignment for the test dataset. These classifiers achieved high performance, with both accuracy and recall exceeding 99% in the validation fold. Classifier performance was evaluated using accuracy, precision, and other classification metrics.

The generated models were compared on the basis of clustering quality metrics (Silhouette score, Davies-Bouldin index, Calinski-Harabasz index), and survival prediction performance (C-index, AIC, log-rank p-value). The robustness of the models was further scrutinized through five-fold stratified cross-validation, bootstrap-based stability assessments (100 iterations) and re-assessed cluster stability (ARI), sample stability, and survival prediction consistency.

The analysis was conducted using Python-based scikit-learn, Matlab, and SciPy tools for clustering and classifier development. The clustering and classification models were implemented using default settings unless otherwise specified.

### Survival Analyses

Survival analysis was conducted using the Kaplan-Meier method to generate overall survival (OS) curves. The log-rank test was used to compare survival distributions between groups. Cases with missing survival status or overall survival time were excluded from analysis. Additionally, Cox proportional hazards regression was performed to assess the effect of hallmark involvement on survival, Survival statistical analyses were performed using SPSS software (version 29.0.0.0 241) and python3. The software was used to perform both the Kaplan-Meier analysis and Cox regression, and the statistical significance of differences in survival distributions was assessed using the log-rank test.

## Results

### Distribution of cancer hallmarks across multiple cancers

The MSK-MET database provides mutation profiles of 25775 tumors of both primary and metastatic origin. One or more hallmark related mutations (HRMs) were identified in 24615 (95.50%) tumours. Invasion/metastasis, escaping cell death, dysdifferentiation and signaling were the most ubiquitous HRMs with on pan-cancer basis (88.1%, 87.8%, 84.4% and 83%; respectively) with ∼2/3 of histopathological subtypes exhibited invasion/metastases and escaping cell death HRMs at frequencies of >80% (56/84 and 54/84; respectively). While the least numbers of HRMs were observed in genes associated with escaping immune response and pathogen response (47.4% and 14.6%, respectively). *TP53* variants were detected in 48.1% of tumors.

HRM patterns varied distinctly across tumor histologies (Supplementary table S3 and suppl Fig 1). Angiogenesis-related HRMs were highly prevalent (>90%) in pancreatic, renal, endometrioid carcinomas, and cutaneous squamous cell carcinoma (SCC) and melanomas, but were observed in only 30.3% of prostatic adenocarcinomas and in <20% of dedifferentiated/well-differentiated liposarcomas and leiomyosarcomas.

Lung adenocarcinoma (LUAD) and lung squamous cell carcinoma (LUSC) exhibited comparable overall HRM frequency profiles, although with significantly different TP53 involvement (47.4% in LUAD vs. 86.6% in LUSC). High HRM prevalence (≥85% of tumors) was observed in LUAD, LUSC, and small cell lung carcinoma across 6/16, 5/16, and 12/16 hallmarks, respectively.

Pancreatic adenocarcinoma and pancreatic neuroendocrine neoplasms had similar tumor mutation burdens (TMB: 3.45 vs. 2.5 muts/MB), yet their HRM profiles diverged notably. Over 90% of pancreatic adenocarcinomas showed HRMs across 8/16 hallmarks, while neuroendocrine neoplasms exhibited no hallmark with >90% HRM prevalence (highest: escape from cell death in 79.6% of tumors).

HPV-related carcinomas (anal and cervical SCC) showed strikingly similar HRM profiles across 15/16 hallmarks, with a significant difference in replicative immortality (Fisher’s exact test, p = 0.04). Oropharyngeal SCC, also associated with HPV, shared HRM similarities across 11/16 hallmarks and differed in 5/16 hallmarks (angiogenesis, energetics, dysdifferentiation, replicative immortality, and TP53 involvement). Overall, these findings suggested that different tumors from the same organ can display markedly different HRM landscapes, and tumors with shared etiology can show significant overlap in the HRM profiles (Fig 1. Table S4).

**Figure 1:**
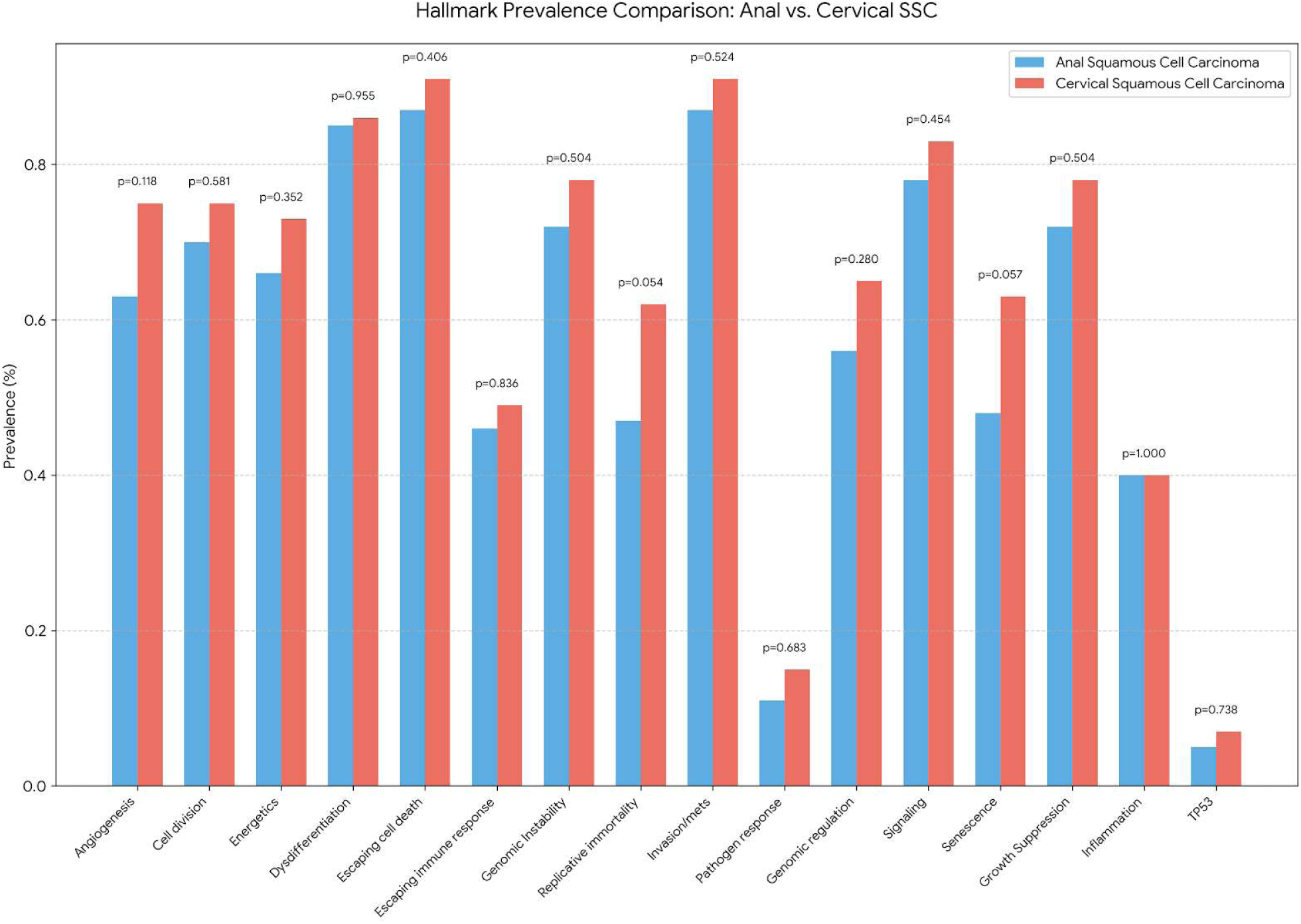
Comparison of hallmark related mutations across HPV related squamous carcinoma.

### Prognostic significance of hallmark involvement

Univariate cox regression analysis showed that HRMs across 14/16 hallmarks exhibited a significant association with overall survival (not statistically significant for cell division and escape immune response). Nearly all the HRMs were associated with inferior survival except for pathogen response which showed marginally better OS (HR: 0.804, p <0.001). Further, the patients without HRM have significantly better overall survival than those with HRM (median OS: 58.97 Vs 40.08 months; log rank p<0.001 and HR: 0.827; 95%CI: 0.764-0.895, p <0.001). Furthe details are described in Table 1.

**Table 1:** The prognostic impact of individual hallmark involvement and results of rule-out dominant gene analysis.

| Hallmark | HR (95% CI);p value | Dominant Gene | Group Cindex | Non Dominant Cindex | Group AIC | Non Dominant AIC |
| --- | --- | --- | --- | --- | --- | --- |
| Angiogenesis | 1.163(1.114-1.214) ;<.001 | *KRAS* | 0.517038 | 0.504667 | 193613.4028 | 193659.6662 |
| Cell division | 1.018(0.978-1.06) ; 0.374 | *APC* | 0.501433 | 0.507162 | 193660.8378 | 193657.5396 |
| Replicative immortality | 1.259(1.21-1.31) ;<.001 | *KRAS* | 0.530477 | 0.50699 | 193531.6564 | 193658.8383 |
| Energetics | 1.306(1.252-1.363) ;<.001 | *KRAS* | 0.53118 | 0.508141 | 193505.5896 | 193656.3556 |
| Dysdifferentiation | 1.129(1.069-1.193) ;<.001 | *KRAS* | 0.507372 | 0.50647 | 193642.3016 | 193651.7785 |
| Escaping immune response | 0.983(0.945-1.022) ; 0.378 | *STK11* | 0.500157 | 0.504257 | 193660.8499 | 193656.783 |
| Escaping cell death | 1.183(1.111-1.259) ;<.001 | *KRAS* | 0.508031 | 0.499836 | 193632.3571 | 193661.568 |
| Genomic Instability | 1.114(1.067-1.163) ;<.001 | *APC* | 0.51113 | 0.516058 | 193637.1557 | 193625.3472 |
| Genomic regulation | 1.089(1.047-1.132) ;<.001 | *STK11* | 0.514307 | 0.511509 | 193643.2804 | 193650.245 |
| Pathogen response | 0.804(0.759-0.852) ;<.001 | *GATA3* | 0.513676 | 0.508349 | 193604.063 | 193635.8263 |
| Invasion/mets | 1.181(1.109-1.257) ;<.001 | *KRAS* | 0.508134 | 0.502365 | 193633.2739 | 193659.568 |
| Signaling | 1.148(1.088-1.21) ;<.001 | *KRAS* | 0.510286 | 0.507082 | 193635.0621 | 193649.1982 |
| Senescence | 1.106(1.064-1.15) ;<.001 | *CDKN2A* | 0.518417 | 0.508165 | 193635.6639 | 193659.6066 |
| Growth Suppression | 1.164(1.114-1.217) ;<.001 | *CDKN2A* | 0.514944 | 0.510027 | 193615.6369 | 193637.3037 |
| Inflammation | 1.25(1.203-1.3) ;<.001 | *KRAS* | 0.530238 | 0.512713 | 193535.2522 | 193644.7845 |
| TP53 | 1.68(1.61-1.74) ;<.001 | *TP53* | NA | NA | NA | NA |

**Table 2:**
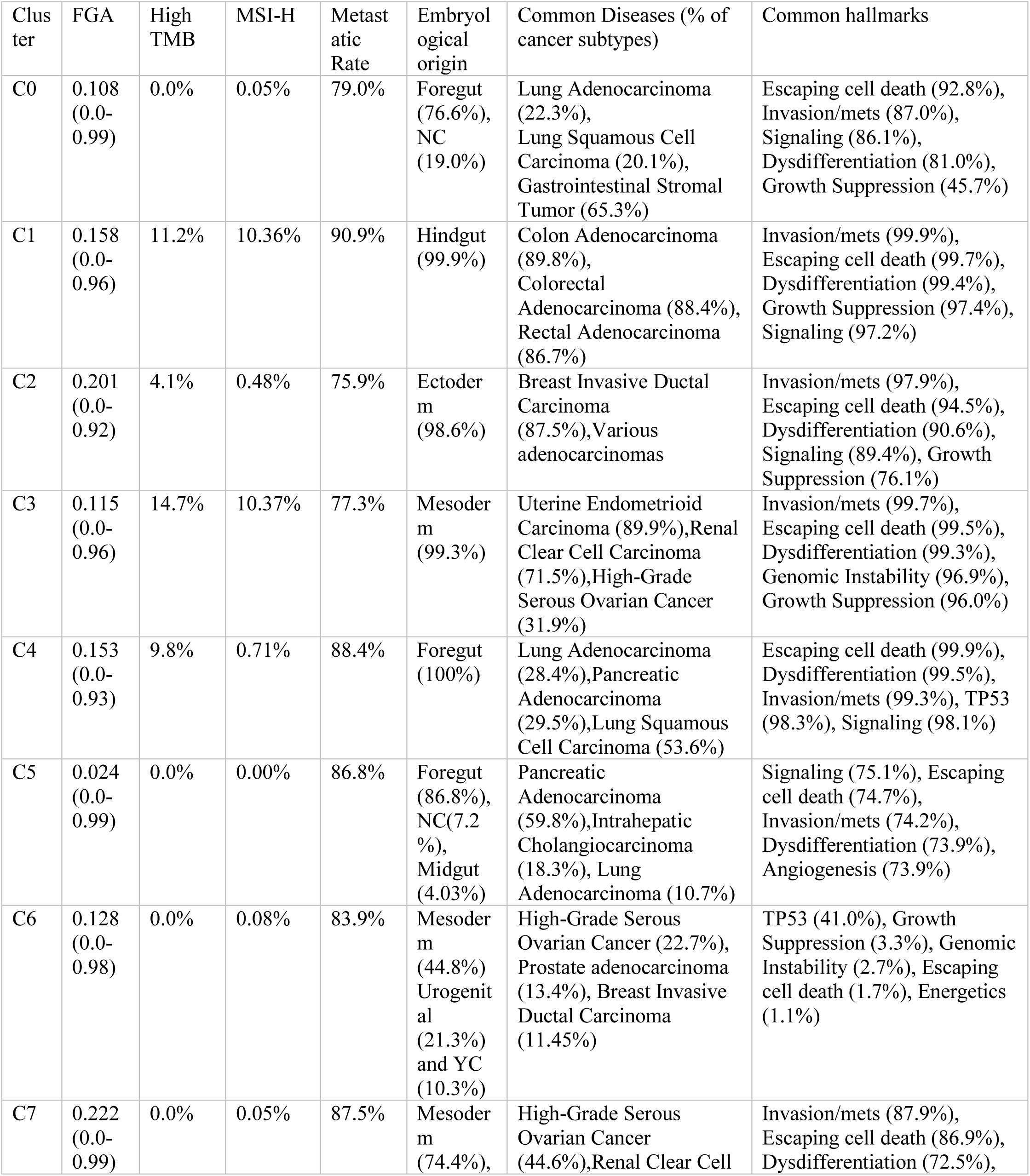
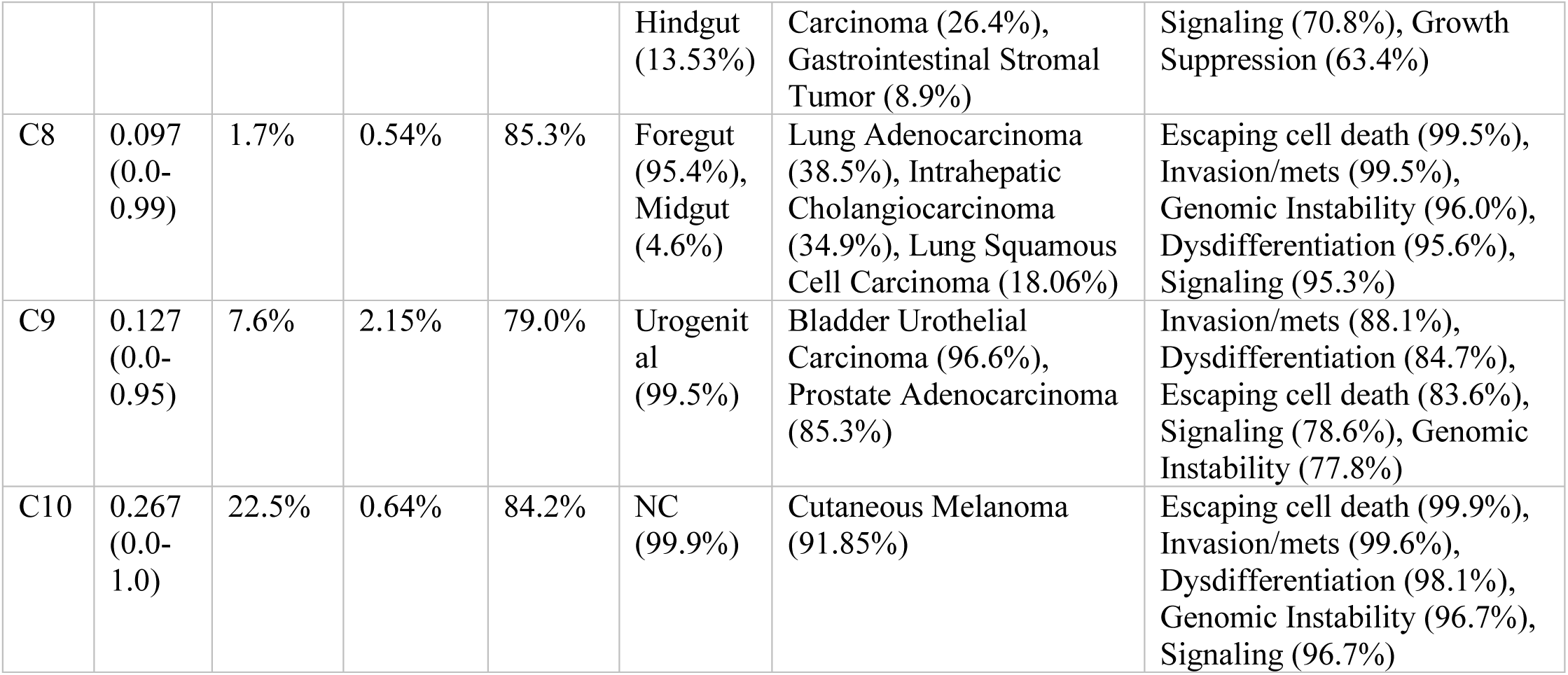
Biological, clinical, embryological profiles of EO/HRM clusters.

Grouping genetic mutations based on shared biological functions can pose analytical challenges, particularly when a single gene—due to high mutation frequency or disproportionately strong prognostic effect—dominates the group’s overall signal. To evaluate the robustness of hallmark gene groups, we conducted a leave-one-out analysis to assess whether their prognostic performance remained intact after excluding the most influential gene. First, the prognostic value of each hallmark gene group was assessed using Cox proportional hazards regression. Univariate Cox regression was applied to each gene to identify the dominant gene within each group—the gene with the highest C-index. Subsequently, differences in the prognostic impact of HRMs, with and without dominant gene, were assessed with changes (delta) in concordance index (C-index) and the Akaike Information Criterion (AIC).

Despite some hallmark groups being strongly influenced by dominant genes (e.g., *KRAS* in multiple hallmarks such as angiogenesis, cellular energetics, and inflammation), we found that most hallmark groups remained predictive of overall survival even after the exclusion of their dominant gene, indicating that their prognostic value is not solely dependent on a single high-impact gene and grouping of genes is an effective strategy (Table 1).

### Development and comparison of biological models

Based on the hypothesis, we developed two models: combined embryological origin (EO)/ HRM and HRM only. The parameters are already described in the methods section (definitions). The tSNE analysis identified 11 distinct clusters for EO/HRM model and 12 clusters for the HRM model. The training dataset was used to develop random Forest based classifiers, and its accuracy was assessed on the validation dataset. Both classifiers showed a very high accuracy and recall (>99%) and data in test dataset was analyzed using classifiers (Table S5).

The results revealed that while HRM only model exhibited slightly better clustering quality (Silhouette score: 0.18 vs. -0.0262, Calinski-Harabasz index: 3158.63 vs. 1306.86); however, EO/HRM exhibited better cluster stability (ARI: 0.7426 vs. 0.7011) and sample stability (0.1989 vs. 0.1798) during cross-validation (bootstrap-based stability assessment using 100 resampling iteration). Among EO/HRM clusters, all 11 showed statistically significant associations with survival, whereas only 8 of 12 HRM-only clusters demonstrated significant survival associations (Suppl table S6). EO/HRM clusters exhibited a higher C-index (0.5917 vs. 0.5649) and lower AIC (192719.17 vs. 193165.76) and this performance advantage was retained during five-fold stratified cross-validation of Cox proportional hazards models (mean C-indices of 0.5907 ± 0.0080 for EO/HRM and 0.5640 ± 0.0054 for HRM-only clusters).

To test generalizability, a re-clustering cross-validation framework was implemented in which clusters were re-derived in each training fold and subsequently applied to the test fold using the nearest-centroid assignment. This more stringent evaluation demonstrated that EO/HRM preserved prognostic stratification with a mean C-index of 0.5742 ± 0.0079, compared with 0.5628 ± 0.0047 for HRM-only. In contrast, HRM-only exhibited higher mean silhouette values (0.2389 vs. 0.1642), indicating tighter geometric separation despite weaker prognostic performance. Collectively, the integration of embryological origin of organ yielded enhanced cluster stability, higher prognostic concordance, and greater reproducibility (Table S6).

### Biological characteristics of Clusters

The EO/HRM clusters exhibited distinct genomic and embryological profiles. Biological properties and distribution across disease subtypes have been described in (Fig 2, Table2, suppl table S7-S10). Cluster C5 was characterized by a genomically quiescent phenotype, with a very low tumor mutational burden (TMB) (median: 2 mutations/Mb; TMB-High: 0%) and minimal genomic instability, as reflected by a low fraction of genome altered (FGA) (median: 0.024). Histologically, C5 was constituted by subgroups of LUAD, PAAD, IHCA and GIST tumours and C5 identified the subgroups with least TMB and FGA among these cancers, confirming quiescent genome across multiple cancer histologies.

**Figure 2:**
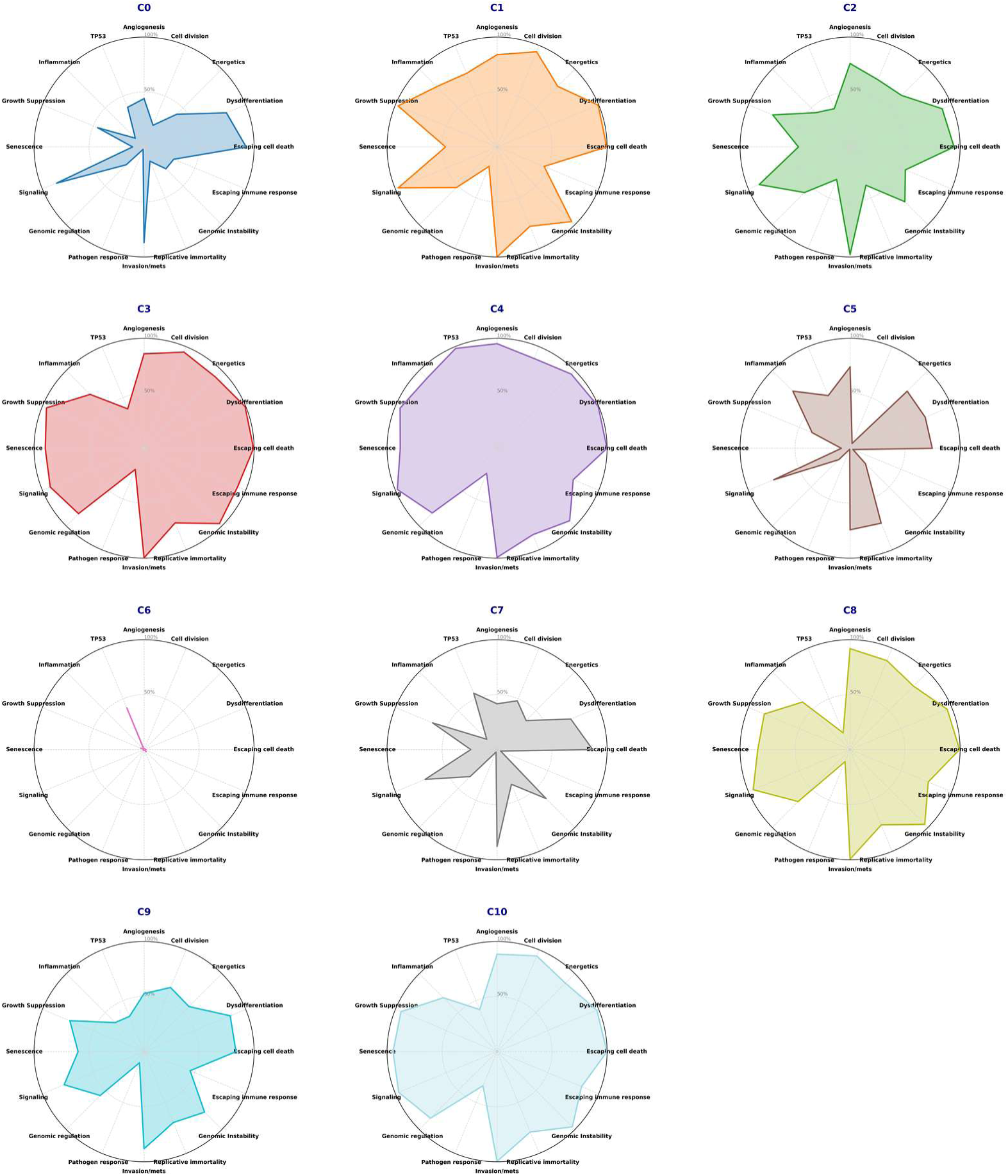
The hallmark related mutation profile of EO/HRM clusters.

In contrast, cluster C10 displayed a highly unstable genomic profile, with the highest TMB (median: 9 mutations/Mb; TMB-High: 22.5%) and the highest FGA (0.267). Embryologically, 86.8% of tumors in C5 originated from the foregut, whereas 99.9% of C10 tumors were derived from the neural crest. The C10 cluster consisted of subgroups of cutaneous melanoma and GIST. While CM is known for high TMB, GIST C10 tumours also exhibited highest TMB among GIST patients (median 2.9 mut/MB; Vs 0-2.6 in other clusters; p<0.001)

Clusters C1 and C3 showed a genome with highest frequencies of microsatellite instability-high (MSI-H) tumors (10.4% each), a comparable high prevalence of TMB-H cases (11.2% and 14.7%), and a moderate level of genomic instability (FGA: 0.158 and 0.115). Cluster C6 exhibited a unique HRM, dominated by TP53 alterations (41.0%) and a near-complete absence or minimal involvement across other hallmarks (<4% in 15 out of 16 categories), suggesting a largely TP53-driven tumor biology. This cluster also had the lowest TMB (median: 0 mutations/Mb; TMB-H: 0%) and MSI-H frequency (0.1%). Histologically, C6 consisted of high grade serous ovarian carcinoma (HGSOC), prostate carcinoma (PRCA), breast invasive ductal carcinoma (IDC), and colon adenocarcinoma (COAD), and subgroup analysis among these cancers showed a similar profile.

### Metastatic patterns and survival analysis

Out of 84 cancer subtypes, we restricted our analysis to 15 major diseases (each with ≥400 patients), as diseases with fewer than 400 patients often consisted of subgroups with insufficient sample sizes for statistical purposes.

Among the 15 major cancer types analyzed, cluster C1 exhibited the highest global prevalence of metastatic disease (90.9%), whereas C3 showed the lowest (71.9%). There were distinct organ-specific predilections. Notably, clusters C1 and C10 showed affinity for lung metastases, while C9 was characterized by a high prevalence of lymph node and bone metastases, with minimal involvement of the liver and lungs (Fig 3, Table S11). Given that pathological diagnosis (cancer subtype) is a well-known predictor for metastatic status and site, we compared the performance of our clustering system against histological classification. Univariate logistic regression models utilizing cluster assignment achieved AUCs (0.68) near equivalent to better performance as compared to histology-based models (0.67). A multivariate model with cancer subtype and cluster as covariates showed that C10, C7, C5 and C4 retained their independent predictive capacity (OR: 2.1, 1.7, 0.78 and 1.6 respectively; p value <0.05, Fig 4). Further, metastasis site specific multivariate logistic regression analysis (Fig 5, Table S13) revealed that C10 was significantly associated with metastases to the lung, liver, and bone (OR: 1.7, 2.0, and 1.8, respectively; p < 0.05 for all), while C4 showed a preferential association with brain metastases (OR: 1.3; p = 0.002). In contrast, clusters C5 and C8 were associated with a reduced risk of lymph node, brain and liver metastases (OR: 0.8; p < 0.05).

**Figure 3:**
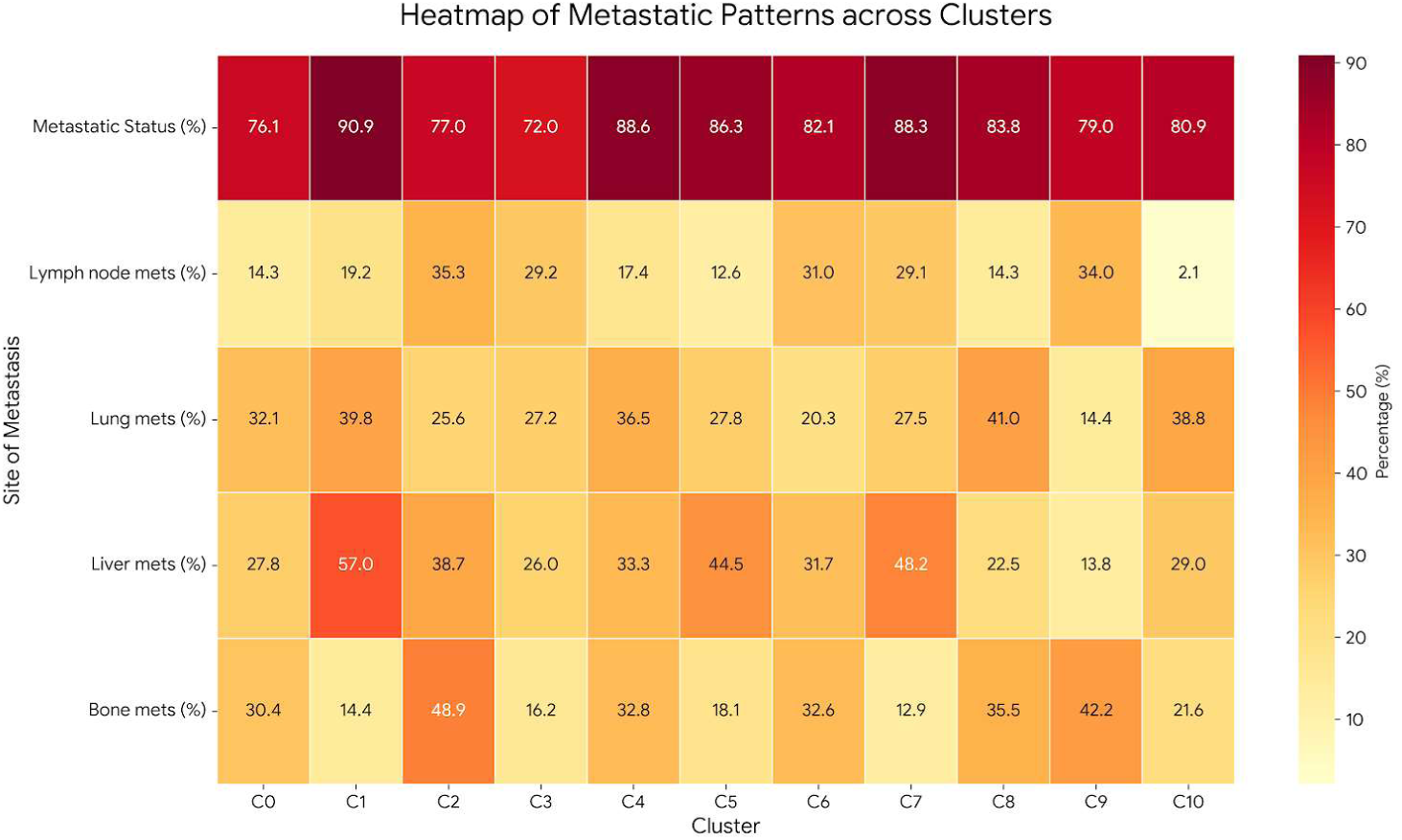
Distribution of clusters across common metastatic sites.

**Figure 4:**
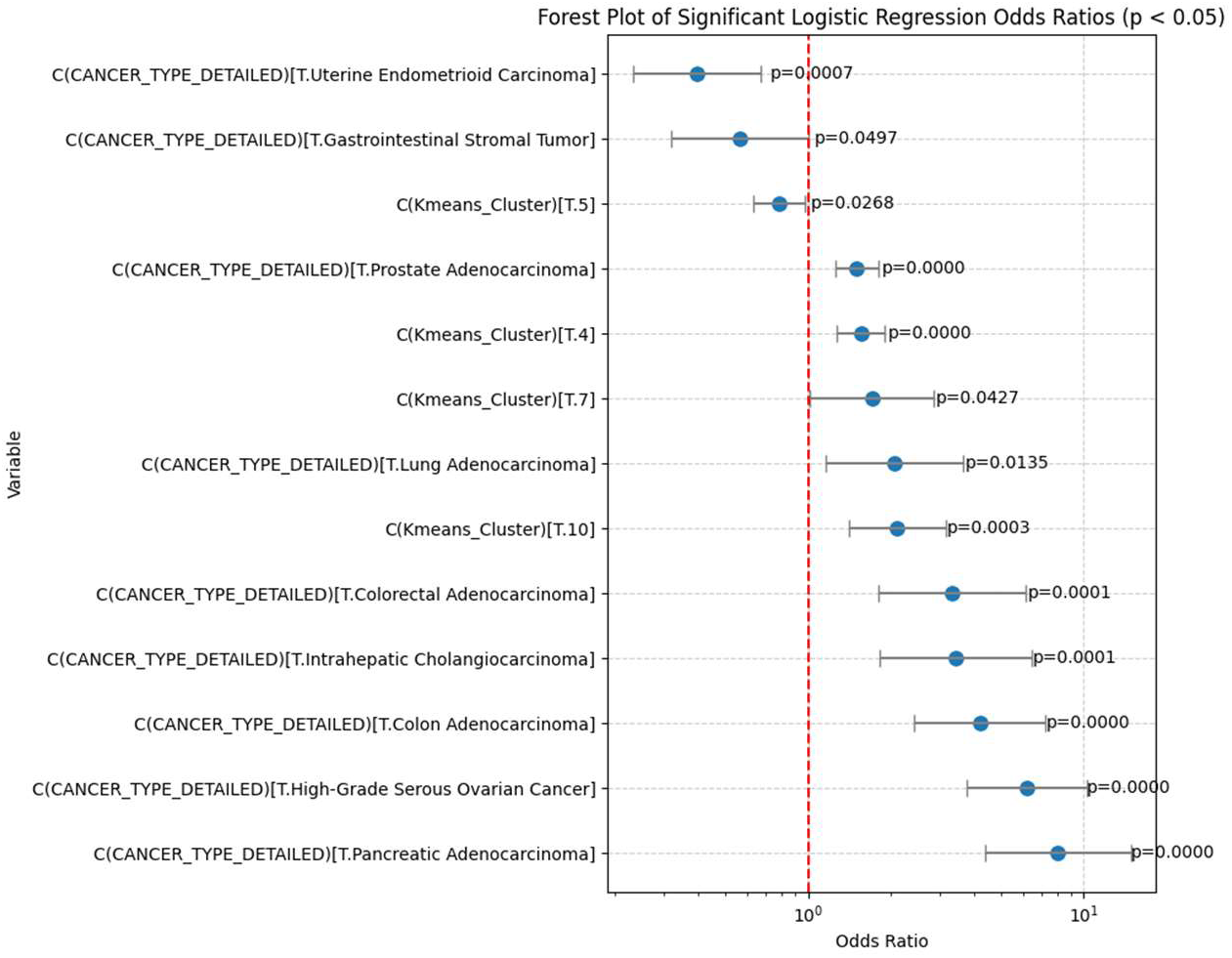
Forest plot depicting results of multivariant regression analysis.

**Figure 5:**
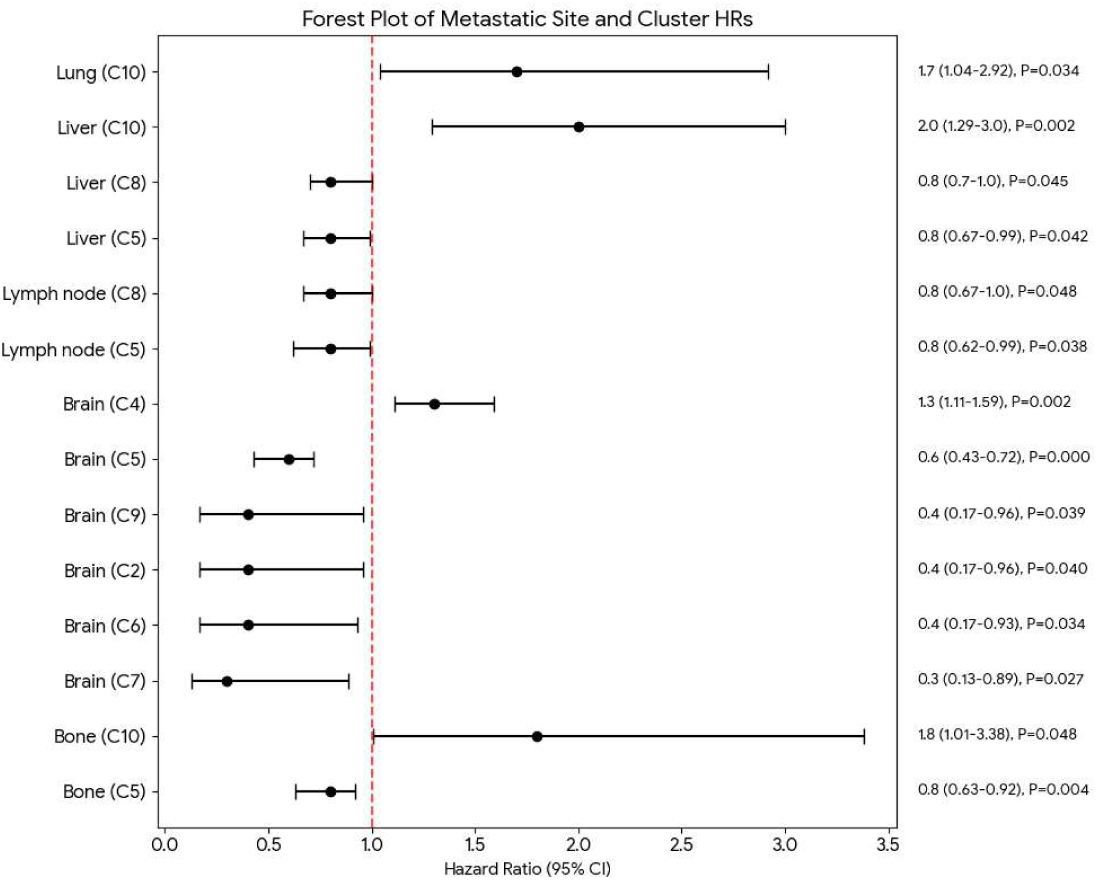
Forest plot describing the odds ratio for metastatic site-specific multivariate analysis.

Further, we identified differences in unique metastatic patterns among individual cancer subtypes (Table 3 and S14). Most notably, cluster C7 demonstrated opposing effects depending on the tissue of origin. While cluster C7 conferred relatively high metastasis risk in high-grade serous ovarian cancer and GIST (97.3% vs 93.8% in C3 and 69.4% vs 51.3% in C0, respectively). Conversely, the same C7 was protective in renal cell clear cell carcinoma, where the prevalence of metastatic disease was lower than other clusters (73% vs 89% in C3). Clusters C0 and C5 were also associated with a lower risk of metastatic disease in lung adenocarcinoma and GIST and exhibited excellent overall survival (C0 & C5-51.9 & 43.2 vs 32.8 to 41.6 months in lung, median OS not reached vs 28.7 months for C7 in GIST). The prognostic signature for C0 and C5 also extended to pancreatic and intrahepatic cholangiocarcinoma. In contrast, lung adenocarcinoma, pancreatic adenocarcinoma, and intrahepatic cholangiocarcinoma patients belonging to cluster C4 had the worst prognosis. Furthermore, a prognostic inversion was observed within prostate carcinoma. While Cluster C0 consistently defined a favorable subgroup in other malignancies, it was associated with the poorest outcomes in the prostate cohort (median overall survival: 29.6 months) and patients belonging to cluster C6 demonstrated significantly superior outcomes (median OS 29.6 months vs not reached, respectively). While a similar trend toward improved survival for Cluster C6 was observed in patients with invasive breast ductal carcinoma and high-grade serous ovarian cancer, these associations did not reach statistical significance.

**Table 3:** median overall survival and prevalence of metastases of EO/HRM clusters across common malignancies.

| Cancer Subtype | Cluster | N | Median OS (Months) | p value | Metastatic Status |  |
| --- | --- | --- | --- | --- | --- | --- |
| Lung Adenocarcinoma | C0 | 908 | 51.9 | <0.001 | 724 (79.74%) | <0.001 |
|  | C4 | 1156 | 32.8 |  | 1012 (87.54%) |  |
|  | C5 | 435 | 43.2 |  | 321 (73.79%) |  |
|  | C8 | 1565 | 41.6 |  | 1292 (82.56%) |  |
| Prostate Adenocarcinoma | C0 | 24 | 29.6 | <0.001 | 18 (75.00%) | 0.7226 |
|  | C2 | 3 | NA |  | 3 (100.00%) |  |
|  | C5 | 1 | NA |  | 1 (100.00%) |  |
|  | C6 | 291 | inf |  | 229 (78.69%) |  |
|  | C9 | 1853 | 72.1 |  | 1511 (81.54%) |  |
| Uterine Endometrioid Carcinoma | C3 | 751 | inf | 0.005 | 424 (56.46%) | 0.5773 |
|  | C6 | 10 | NA |  | 6 (60.00%) |  |
|  | C7 | 74 | inf |  | 48 (64.86%) |  |
| High-Grade Serous Ovarian Cancer | C2 | 7 | NA | 0.173 | 5 (71.43%) | 0.0043 |
|  | C3 | 318 | 39.1 |  | 295 (92.77%) |  |
|  | C6 | 226 | 35.2 |  | 215 (95.13%) |  |
|  | C7 | 445 | 35.1 |  | 433 (97.30%) |  |
|  | C10 | 1 | NA |  | 1 (100.00%) |  |
| Pancreatic Adenocarcinoma | C0 | 57 | 26.6 | <0.001 | 55 (96.49%) | 0.737 |
|  | C4 | 524 | 13.1 |  | 496 (94.66%) |  |
|  | C5 | 1063 | 14.9 |  | 991 (93.23%) |  |
|  | C8 | 135 | 19.4 |  | 127 (94.07%) |  |
| Cutaneous Melanoma | C0 | 40 | inf | 0.048 | 28 (70.00%) | 0.2828 |
|  | C5 | 8 | NA |  | 6 (75.00%) |  |
|  | C7 | 9 | NA |  | 6 (66.67%) |  |
|  | C10 | 642 | inf |  | 527 (82.09%) |  |
| Bladder Urothelial Carcinoma | C0 | 5 | NA | 0.296 | 3 (60.00%) | 0.4787 |
|  | C6 | 28 | inf |  | 24 (85.71%) |  |
|  | C9 | 928 | inf |  | 687 (74.03%) |  |
| Rectal Adenocarcinoma | C1 | 677 | 49.9 | 0.126 | 564 (83.31%) | 0.9698 |
|  | C2 | 1 | NA |  | 1 (100.00%) |  |
|  | C6 | 22 | 22 |  | 19 (86.36%) |  |
|  | C7 | 81 | 35.3 |  | 66 (81.48%) |  |
| Colorectal Adenocarcinoma | C1 | 402 | 57 | 0.371 | 368 (91.54%) | 0.9969 |
|  | C6 | 11 | NA |  | 10 (90.91%) |  |
|  | C7 | 41 | 32.7 |  | 38 (92.68%) |  |
|  | C9 | 1 | NA |  | 1 (100.00%) |  |
| Breast Invasive Ductal Carcinoma | C0 | 2 | NA | 0.853 | 2 (100.00%) | 0.4243 |
|  | C2 | 1957 | 57.9 |  | 1507 (77.01%) |  |
|  | C5 | 10 | NA |  | 5 (50.00%) |  |
|  | C6 | 256 | 72.5 |  | 189 (73.83%) |  |
|  | C7 | 8 | NA |  | 6 (75.00%) |  |
|  | C9 | 3 | NA |  | 2 (66.67%) |  |
| Lung Squamous Cell Carcinoma | C0 | 108 | 22.7 | 0.67 | 82 (75.93%) | 0.4501 |
|  | C4 | 288 | 20.6 |  | 236 (81.94%) |  |
|  | C5 | 44 | 18.1 |  | 34 (77.27%) |  |
|  | C8 | 97 | 22.3 |  | 83 (85.57%) |  |
| Intrahepatic Cholangiocarcinoma | C0 | 159 | 16.1 | 0.024 | 145 (91.19%) | 0.3209 |
|  | C4 | 31 | 7.2 |  | 27 (87.10%) |  |
|  | C5 | 75 | 14.1 |  | 61 (81.33%) |  |
|  | C8 | 142 | 14.6 |  | 123 (86.62%) |  |
| Renal Clear Cell Carcinoma | C3 | 301 | 59.5 | 0.385 | 268 (89.04%) | <0.001 |
|  | C6 | 9 | NA |  | 3 (33.33%) |  |
|  | C7 | 111 | inf |  | 81 (72.97%) |  |
| Gastrointestinal Stromal Tumor | C0 | 263 | inf | 0.006 | 135 (51.33%) | 0.0248 |
|  | C3 | 1 | NA |  | 0 (0.00%) |  |
|  | C5 | 25 | inf |  | 14 (56.00%) |  |
|  | C7 | 36 | 28.9 |  | 25 (69.44%) |  |
|  | C10 | 78 | inf |  | 55 (70.51%) |  |
| Colon Adenocarcinoma- Low TMB | C0 | 1 | NA | 0.5 | 1 (100.00%) | 0.9311 |
|  | C1 | 1797 | 35.5 |  | 1703 (94.77%) |  |
|  | C2 | 1 | NA |  | 1 (100.00%) |  |
|  | C6 | 45 | 35.2 |  | 42 (93.33%) |  |
|  | C7 | 189 | 52.4 |  | 175 (92.59%) |  |
|  | C9 | 1 | NA |  | 1 (100.00%) |  |

## Discussion

Recent advances in genomic sequencing and epigenomic profiling (such as methylation landscapes) have provided significant insights into the molecular basis of cancer. While this information has refined precision medicine, current models based solely on individual alterations often fail to provide a comprehensive or reproducible system for predicting the natural history of disease or the exact response to targeted therapies. Consequently, there is a critical need for integrated models that represent the complex, multifactorial nature of the neoplastic cell.

Dynamic systems theory provides a robust framework for this complexity by predicting the existence of attractor states—stable equilibria toward which a system gravitates despite various influencing factors. In this context, we conceptualize the cancer cell as a complex system defined by the interplay of somatic genomic variants, dysregulated epigenomics, and primitive cellular programs. We hypothesize that the embryological cell of origin (EO) serves as a proxy for the inherent “epigenetic topography,” while hallmark-related mutations (HRM) represent “topographical perturbations” that result in the formation of alternative “pathological basins” or valleys.

To identify potential cancer attractor states, we employed a large-scale genomic database and analyzed it using a novel hallmark-based system. Our analysis demonstrated that this hallmark-based interpretation of genomic variants reveals distinct subgroups with unique biological profiles and clinical behaviors across multiple histologies. Critically, these clusters represent specific co-occurrences of acquired neoplastic properties developed in the context of primitive cellular programming. The robustness of this system was confirmed through comprehensive validation, including bootstrap resampling, five-fold cross-validation, and re-clustering procedures, ensuring that these patterns are highly stable and reproducible.

To demonstrate effectiveness of including inherent cellular program in the model, we compared two models: HRM-only and combining embryological origin with HRM (EO/HRM). Both models identified biologically distinct subsets; but EO/HRM model demonstrated superior stability, reproducibility, and marginally improved survival prediction. Overall, these results indicated that incorporating inherent cellular programing (GRNs) of cell of origin, even via a coarse marker such as embryonic origin, enhances the identification of stable, prognostically relevant patterns at both pan-cancer and cancer subtypes levels.

The clustering model identified disease subgroups with unique clinical behavior and disease biology. Different clusters exhibited variable risks for metastases and distinct predilection for specific metastatic organs, a pattern which persisted even after correcting known determinants like histological subtype. While clusters C1, C3, C4, C10 showed a high proportion of TMB-H cancers, only C1 and C3 exhibited concordant high prevalence of MSI-H, thus highlighting a non-MSI driver mechanism in C4 and C10. Among C6 cancers, *TP53* mutations were seen in nearly half of patients and showed near total absence of any other hallmark, indicating a unique underlying pathophysiological mechanism in these cancers. This subgroup consisted mainly of high grade serous ovarian carcinoma and substantial subsets of breast invasive ductal carcinoma and prostatic carcinoma, where it exhibited a favorable prognosis later. Further, survival analysis of clusters showed contradictory trends across multiple malignancies e.g. C0 was generally associated with favorable prognosis in LUAD, PAAD, IHCA but poor prognosis in PRCA. It highlighted limitations of prognostic interpretation of genomic data in a tissue agnostic manner.

Hallmark-based groupings offer a framework for personalized precision medicine strategies. Angiogenesis-associated variants were highly prevalent in pancreatic, renal, and melanoma tumors and correlated with poor prognosis, underscoring their contribution to oncogenesis and potential as biomarkers for VEGF/angiogenesis inhibitor therapy(17,18). Conversely, pathogen-response variants were associated with improved survival, suggesting enhanced immune responsiveness and warranting evaluation as predictive biomarkers for immunotherapy, like MSI-H status. In contrast, variants promoting immune evasion may indicate resistance to immune checkpoint inhibitors. Distinct metastatic profiles across clusters provide opportunities for personalized, site-specific surveillance—for example, Cluster C10 exhibited increased risk for bone, liver, and lung metastases, whereas Cluster C5 showed minimal metastatic potential. Tumor-agnostic therapies, targeting genomic biomarkers irrespective of histology, have demonstrated variable efficacy across malignancies; the EO/HRM model highlights the influence of cell-of-origin in shaping disease behavior. This finding supports the development of integrated models incorporating both genomic alterations and cell of origin as a predictive biomarker and identifying patient subgroups most likely to benefit from tissue agnostic approaches.

The application of hallmark and embryonic expression profile-based prognostications has been explored in separate studies. Nagy et al. demonstrated that RNA-seq expression profiling of hallmark-related genes, particularly those associated with genomic instability and invasion/metastases, yields statistically significant prognostic data (8). Similarly, Ben-Porath et al. found that poorly differentiated carcinomas exhibit gene expression profiles overlapping with embryonic stem cells (notably the Yamanaka factors: *Nanog, Oct4, Sox2,* and *c-Myc*), which correlates with poor outcomes in glioma and bladder carcinoma (11). More recently, Barcelos et al. identified key Gene Regulatory Networks (GRNs) in glial neoplasms, proving that the expression profiles of genes within these networks offer relevant prognostic value. However, these studies rely on RNA-seq data, which is not feasible in routine clinical practice, thereby limiting the clinical utility of such signatures. Hence, the present study focuses on categorizing genomic variants, rather than expression profiles, by their relationship to cancer hallmarks with a central premise is that genomic alterations are not stochastic and they emerge systematically, influenced by the cell’s specific GRN, to facilitate the acquisition of neoplastic properties

This study had several limitations. First, copy number variations and non-fusion structural variants were excluded due to interpretation challenges, and it may have influenced hallmark profiles. Second, hallmark annotations assumed uniform direction i.e.; mutation will either cause promotion of oncogenic or inhibition of cancer suppressive hallmarks. Also, hallmark association was measured as a binary parameter, which may not account for mutation burden in individual hallmarks. Third, embryological origin was inferred from organ-level classification, which may not fully capture the inherent cellular programs of cancer cells. Additionally, the MSK-MET dataset predominantly represents advanced-stage and metastatic cancers, limiting generalizability to early-stage disease and survival models could not be adjusted for other known parameters including age, performance status, and treatment response. While internal validation using cross-validation and bootstrap methods demonstrated robustness, external validation on independent cohorts is necessary to confirm reproducibility. Finally, this analysis is computational and lacks experimental validation of hallmark-driven clusters, restricting mechanistic interpretation. However, despite the limitations, this study did show that hallmark-based gene grouping can be a useful tool for interpretation of the complex mutation landscape of cancers and can provide clinically relevant information.

## Conclusion

The concepts of hallmarks of cancer and attractor states provide a robust framework for understanding complex interplay between modulators of cancer cells and development of better predictive biomarkers. In this study, we demonstrated that annotating somatic variants based on hallmark provided relevant prognostic information. Further, integrating embryological origin with hallmark-related mutations enables the unsupervised identification of distinct and stable attractor states across human cancers. The EO/HRM system provides a robust model with the ability to identify clinically, biologically and prognostically distinct subgroups with unique genomic architectures, metastatic patterns, and survival trends. However, external validation and experimental studies are needed to confirm biological mechanism and clinical applicability. Collectively, this study describes a novel and practical strategy for functional interpretation of complex mutational landscapes, genomic risk stratification with a potential for further refinement of genomic guided therapies.

